# Pattern of early human-to-human transmission of Wuhan 2019-nCoV

**DOI:** 10.1101/2020.01.23.917351

**Authors:** Julien Riou, Christian L. Althaus

## Abstract

On December 31, 2019, the World Health Organization was notified about a cluster of pneumonia of unknown aetiology in the city of Wuhan, China. Chinese authorities later identified a new coronavirus (2019-nCoV) as the causative agent of the outbreak. As of January 23, 2020, 655 cases have been confirmed in China and several other countries. Understanding the transmission characteristics and the potential for sustained human-to-human transmission of 2019-nCoV is critically important for coordinating current screening and containment strategies, and determining whether the outbreak constitutes a public health emergency of international concern (PHEIC). We performed stochastic simulations of early outbreak trajectories that are consistent with the epidemiological findings to date. We found the basic reproduction number, *R*_0_, to be around 2.2 (90% high density interval 1.4—3.8), indicating the potential for sustained human-to-human transmission. Transmission characteristics appear to be of a similar magnitude to severe acute respiratory syndrome-related coronavirus (SARS-CoV) and the 1918 pandemic influenza. These findings underline the importance of heightened screening, surveillance and control efforts, particularly at airports and other travel hubs, in order to prevent further international spread of 2019-nCoV.

## Introduction

On December 31, 2019, the World Health Organization (WHO) was alerted about a cluster of pneumonia of unknown aetiology in the city of Wuhan, China[1]. Only a few days later, Chinese authorities identified and characterised a novel coronavirus (2019-nCoV) as the causative agent of the outbreak [2]. The outbreak apparently started from a single or multiple zoonotic transmission events at a wet market in Wuhan, and has since resulted in 655 confirmed cases in China and several other countries by January 23, 2020 [3]. At this early stage of the outbreak, it is critically important to gain a better understanding of the transmission pattern and the potential for sustained human-to-human transmission of 2019-nCoV. Information on the transmission characteristics will help coordinate current screening and containment strategies, support decision making on whether the outbreak constitutes a public health emergency of international concern (PHEIC), and is necessary for anticipating the risk of pandemic spread of 2019-nCoV.

Two key properties will determine further spread of 2019-nCoV. First, the basic reproduction number *R*_0_ describes the average number of secondary cases generated by an infectious index case during the early phase of the outbreak. If*R*_0_ is above the critical threshold of 1, continuous human-to-human transmission with sustained transmission chains will occur. Second, the individual variation in the number of secondary cases provides further information about the early outbreak dynamics and the expected number of superspreading events [4, 5, 6].

In order to better understand the early transmission pattern of 2019-nCoV, we performed stochastic simulations of early outbreak trajectories that are consistent with the epidemiological findings to date.

## Methods

We performed stochastic simulations of the first few generations of human-to-human transmission of 2019-nCoV. Simulations were initialized with one index case. For each primary case, we generated secondary cases according to a negative-binomial offspring distribution with mean *R*_0_ and dispersion *k* [4, 5]. The dispersion parameter *k* can be interpreted as a measure of the probability of superspreading events (the lower the value of k, the higher the probability of superspreading). The generation time interval *D* was assumed to be gamma-distributed with a shape parameter of 2, and a mean that varied between 7 and 14 days. We explored a wide range of parameter combinations (Table 1) and ran 1,000 stochastic simulations for each individual combination. This corresponds to a total of 3.52 million one-index-case simulations that were run on UBELIX (http://www.id.unibe.ch/hpc), the high performance computing cluster at the University of Bern.

**Table 1:** Parameter ranges for stochastic simulations of outbreak trajectories.

| Parameter | Description | Range |
| --- | --- | --- |
| $R_0$ | Basic reproduction number | [0.8 – 5.0] |
| $k$ | Dispersion parameter | [0.01 – 10] |
| $D$ | Generation time interval | [7 – 14] |
| $n$ | Initial number of index cases | [1 – 50] |
| $T$ | Date of zoonotic transmission | [20 Nov 2019 – 4 Dec 2019] |

In a second step, we accounted for the uncertainty regarding the number of index cases *n* and the date *T* of the initial zoonotic animal-to-human transmissions at the wet market in Wuhan. An epidemic with several index cases can be considered as the sum of several independent epidemics with one index case each. We sampled (with replacement) *n* of the one-index-case epidemics, sampled a date of onset for each index case, and summed the epidemic curves together. The sampling of the date of onset was done uniformly from a two-week interval around November 27, 2019, in coherence with early phylogenetic analyses of 11 2019-nCoV genomes [7]. This step was repeated 4,800 times for each combination of *R*_0_ (22 points) and *k* (20 points) for a total of 2,112,000 full epidemics simulated that included the uncertainty on *D, n* and *T*. Finally, we calculated the proportion of stochastic simulations that reached a total number of infected cases within the interval [1000, 9700] by January 18, 2020, as estimated by Imai and colleagues [8]. In a process related to Approximate Bayesian Computation (ABC), the parameter value combinations that led to simulations within that interval were treated as approximations to the posterior distributions of the parameters with uniform prior distributions. Model simulations and analyses were performed in the R software for statistical computing [9]. Code files are available on https://github.com/jriou/wcov.

## Results

In order to reach between 1,000 and 9,700 infected cases by January 18, 2020, the early human-to-human transmission of 2019-nCoV are characterized by values of *R*_0_ around around 2.2 (90% high density interval 1.4-3.8) (figure 1). The observed data at this point is compatible with a large range of values for the dispersion parameter *k* (median 0.54, 90% high density interval 0.014-6.95). However, our simulations suggest that very low values of *k*, corresponding to a large probability of superspreading events, are less likely. These estimates incorporate the uncertainty on the current total epidemic size (as of January 23, 2020) and on the date and scale of the initial zoonotic event (figure 2).

**Figure 1:**
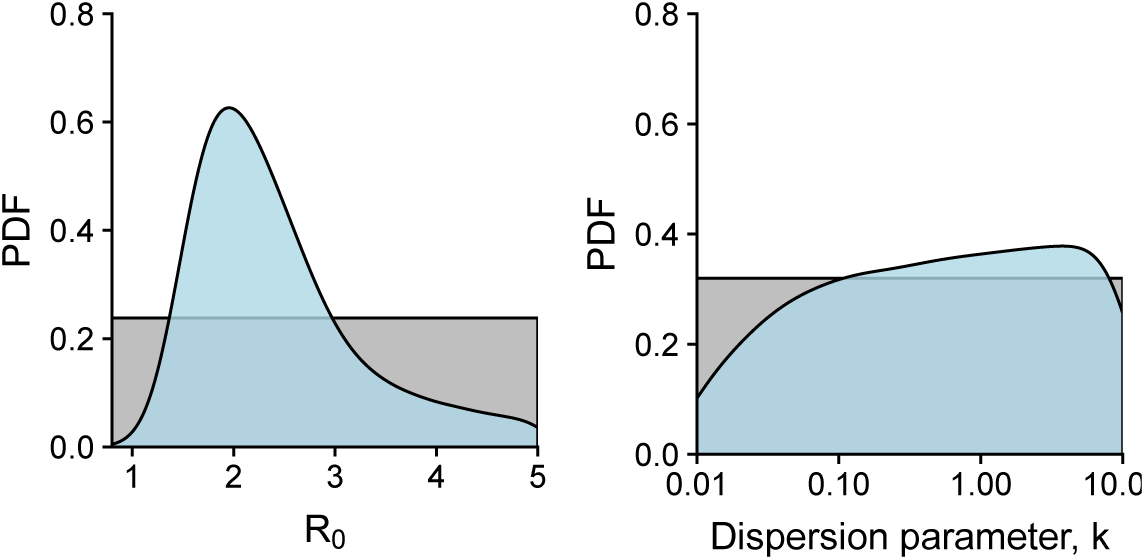
Values of *R*_0_ and *k* most compatible with epidemic data available on 2019-nCoV as of January 23, 2020 (in blue). The basic reproduction number *R*_0_ quantifies human-to-human transmission. The dispersion parameter *k* quantifies the risk of a superspreading event (lower values of *k* are linked to a higher probability of superspreading).

**Figure 2:**
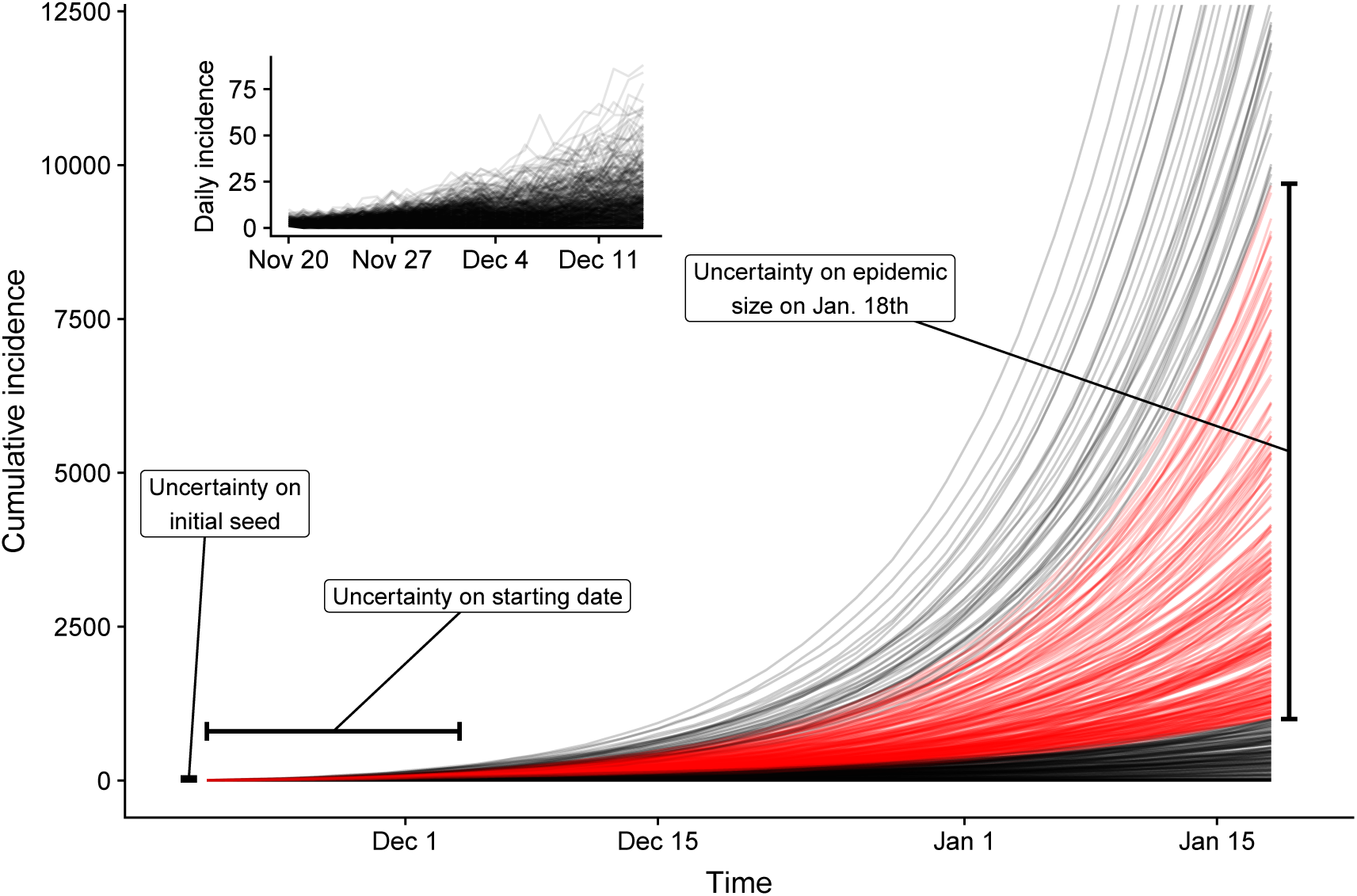
Illustration of the simulation strategy. The lines represent the cumulative incidence of 100 simulations with *R*_0_ = 1.8 and *k =* 1.13. The other parameters are left to vary according to table 1. Among these simulated epidemics, 54.3% led to a cumulative incidence between 1000 and 9700 on January 18, 2020 (in red, as in figure 2).

Comparison with other emerging viruses in the past allows to put in perspective the available information regarding the transmission patterns of 2019-nCoV. Our estimates of *R*_0_ and *k* are more similar to previous estimates focusing on early human-to-human transmission of SARS-CoV in Beijing and Singapore[4] than of MERS-CoV[6] (figure 3). Our estimates for 2019-nCoV are also in line with those of 1918 Influenza [10].

**Figure 3:**
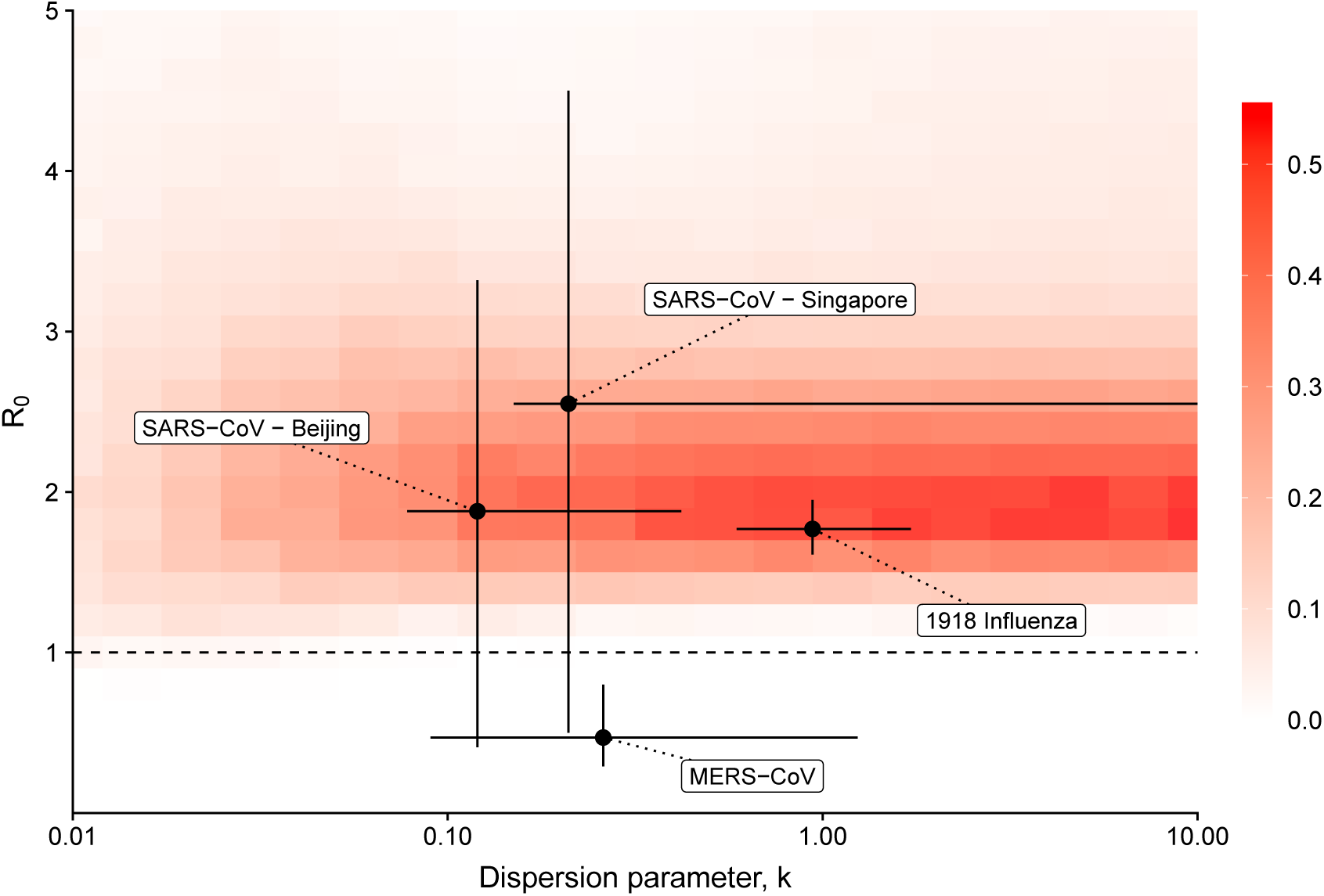
Proportion of simulated epidemics that lead to a cumulative incidence between 1000 and 9700 on January 18, 2020. This can be interpreted as the combinations of *R*_0_ and *k* values most compatible with epidemic data available on 2019-nCoV as of January 23, 2020. As a comparison, we show the estimates of *R*_0_ and *k* for the early human-to-human transmission of SARS-CoV in Singapore and Beijing, and of 1918 Influenza [4, 10, 6].

## Discussion

After SARS-CoV in 2002 and of MERS-CoV in 2012, the emergence of 2019-nCoV in Wuhan, China raises serious concerns about an emerging global health threat. The WHO International Health Regulations Emergency Committee concluded on January 22, 2020 that the lack of information about the patterns of transmission of the disease limited their ability to declare a PHEIC [11]. The increasing research activity on 2019-nCoV has led to estimates of animal-to-human transmission [12], estimates of epidemic size using air travel data [8, 13], and a better characterisation of the epidemiological and virological characteristics of the virus [14, 2]. Our study is the first to focus on the quantification of early human-to-human transmission of 2019-nCoV. The observed patterns of transmission confirm that the current situation has a much closer relationship to what was observed during the early stages of SARS-CoV transmission in Beijing and Singapore rather than during the emergence of MERS-CoV in the Middle-East [4, 6]. Although very different virological characteristics limit the relevance of such a comparison, the transmission pattern of nCoV-2019 is also similar to those of 1918 pandemic influenza [10].

The scarcity of available data, especially on case counts by date of disease onset as well as contact tracing, greatly limits the precision of our estimates and does not yet allow for reliable forecasts of epidemic spread. While based on few data points, our analysis can still shed light on the early transmission pattern of 2019-nCoV. First, our estimate of R0 suggests that the disease has the potential for sustained human-to-human transmission. This implies that effective prevention and control measures will be crucial at this stage to limit further spread of the virus. The recent shutdown of several cities including Wuhan shows that Chinese authorities are aware of the potential magnitude of this outbreak. Second, the risk of superspreading events could be similar to SARS-CoV and MERS-CoV, but is unlikely to be higher. This has important implications for international travel, as superspreading increases the risk of large infection clusters in distant countries originating from one or a few unidentified imported cases. The implementation of control measures in hospital settings, especially emergency rooms, will also be of prime importance, as has been shown by the examples of MERS-CoV in South Korea [15] and in Saudi Arabia [16]. Third, our simulations are also consistent with lower levels of superspreading that are more similar to 1918 pandemic influenza. In this case, we would expect less explosive outbreaks resulting from single cases but a higher risk of sustained transmission chains [4].

Our analysis, while limited due to the scarcity of data, has two important strengths. First, it is based on the simulation of a wide range of possibilities regarding epidemic parameters and allows for the full propagation of the many remaining uncertainties regarding 2019-nCoV and the situation in Wuhan: the size of the initial zoonotic event at the wet market, the date(s) of the initial animal-to-human transmission event(s) and the generation time interval. While accounting for all these uncertainties, our analysis provides a reliable summary of the current state of knowledge about the human-to-human transmissibility of 2019-nCoV. Second, its focus on the possibility of superspreading events by using negative-binomial offspring distributions is very important in the context of emerging coronaviruses [4, 5]. While our estimate of *k* remains very imprecise, the simulations suggest that very low values of *k* (< 0.1) are less likely than higher values (> 1) that correspond to a more homogeneous transmission pattern. However, values of k in the range of 0.1-0.2 are still compatible with the occurrence of large superspreading events, especially in hospital settings [15, 16].

Our analysis suggests that the early pattern of human-to-human transmission 2019-nCoV is reminiscent of SARS-CoV emergence in 2002. International collaboration and coordination will be crucial in order to contain the spread of 2019-nCoV. At this stage, particular attention should should be given on the prevention of superspreading events, while the rapid establishment of sustained transmission chains from single cases cannot be ruled out. The previous experience with SARS-CoV has shown that established practices of infection control can stop such an epidemic. Given the uncertainty around the case fatality rate, our findings highlight the importance of heightened screening, surveillance and control efforts, particularly at airports and other travel hubs, in order to prevent further international spread of 2019-nCoV.

## Acknowledgements

JR is funded by the Swiss National Science Foundation (grant 174281).

## Authors’ contributions

JR and CLA designed the study, JR performed model simulations, JR and CLA analyzed and interpreted the results and wrote the manuscript.

